# Goodness-of-fit filtering in classical metric multidimensional scaling with large datasets

**DOI:** 10.1101/708339

**Authors:** Jan Graffelman

**Affiliations:** Department of Statistics and Operations Research, Universitat Politècnica de Catalunya; Department of Biostatistics, University of Washington

**Keywords:** Plot brushing, outlier, attractor point, eigenvalue, Manhattan distance, allele sharing distance

## Abstract

Metric multidimensional scaling (MDS) is a widely used multivariate method with applications in almost all scientific disciplines. Eigenvalues obtained in the analysis are usually reported in order to calculate the overall goodness-of-fit of the distance matrix. In this paper, we refine MDS goodness-of-fit calculations, proposing additional point and pairwise good-ness-of-fit statistics that can be used to filter poorly represented observations in MDS maps. The proposed statistics are especially relevant for large data sets that contain outliers, with typically many poorly fitted observations, and are helpful for improving MDS output and emphasising the most important features of the dataset. Several goodness-of-fit statistics are considered, and both Euclidean and non-Euclidean distance matrices are considered. Some examples with data from demographic, genetic and geographic studies are shown.

## 1 Introduction

Multidimensional scaling (MDS) is a versatile multivariate technique that has found application in many branches of science. The goal of the method is to construct a configuration of points in a low-dimensional space, such that inter-point distances in this configuration approximate the entries of a given distance matrix (Torgerson, 1958; Mardia et al., 1979; Johnson and Wichern, 2002). In this era of large datasets, MDS applications can involve distance matrices withhundreds or thousands of rows, leading to dense maps with many points. Not all observations are equally well represented in the map, and some observations may have a poor goodness-of-fit, in the sense that their interpoint distances with other observations poorly approximate the corresponding entries in the original distance matrix. In classical metric MDS, eigenvalues (which represent variance accounted for (Carroll and Chang, 1970)) are used to assess the overall goodness-of-fit of the map, but goodness-of-fit statistics at the level of individual points or for a single pair of observations are lacking. The main idea of this paper is to develop point and pairwise goodness-of-fit statistics, in order to use them for *plot brushing*: by not plotting the poorly represented points the final map will be less dense, and the main features of the data set are better emphasized. At the same time, brushing also avoids misinterpretations based on poorly represented observations. In applications, one is often tempted to interpret all points in the same way, as if all points were equally well represented, but in practice, large differences in goodness-of-fit among observations do often exist. The remainder of this paper is structured as follows. In Section 2 we briefly summarize the theory of classical MDS and develop point and pairwise goodness-of-fit statistics. In Section 3 we apply our statistics to examples with datasets taken from demography, genetics and geography. Section 4 finishes the article with a discussion.

## 2 Theory

We summarize classical metric MDS and establish our notation in Section 2.1. We address point and pairwise goodness-of-fit measures for (pairs of) observations for Euclidean distance matrices in Section 2.2, and discuss analogous measuresfor non-Euclidean distance matrices in Section 2.3.

### 2.1 Classical metric MDS

Classical metric MDS, also known as *classical scaling* or *principal coordinate analysis* (PCO), is a standard topic in courses on multivariate analysis. Text books on multivariate analysis usually dedicate a chapter to this method (Mardia et al., 1979; Johnson and Wichern, 2002). The books of Borg and Groenen (2005) and Cox and Cox (2001) are entirely dedicated to the topic. In this paper we confine ourselves to principal coordinate analysis, which is popular in many fields. Non-metric methods (Kruskal, 1964) and metric methods based on iterative stress minimization (De Leeuw and Mair, 2009) are not considered here. In PCO (Gower, 1966), the approximation of the distance matrix is indirect, via the scalar product matrix **B**, which is obtained by scaling and double-centring the matrix of squared distances

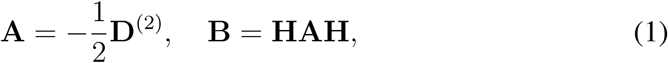

where **D**^(2)^ refers to the *n* × *n* distance matrix with squared entries, and **H** is the centring matrix 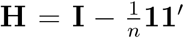. The scalar product matrix **B** is decomposed by the spectral decomposition

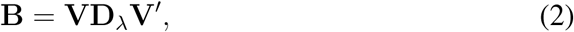

where **V** is the *n* × *n* matrix of orthogonal eigenvectors (**V′V** = **I**_*n*_), and **D**_*λ*_ is a *n* × *n* diagonal matrix containing the eigenvalues of **B** in non-increasing order of magnitude (*λ*_1_ ≥ *λ*_2_ ≥ *…* ≥ *λ*_*n*_). The coordinates of the observations in theMDS map are obtained by

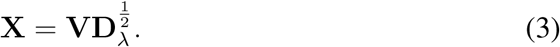

Typically, a two-dimensional representation is made by only using the first two eigenvectors and eigenvalues. The Euclidean distances between the rows of **X**, which we represent by 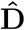, are then used to approximate the original entries of **D**.

We will use the following notation in our goodness-of-fit calculations, and let *g* indicate the overall goodness-of-fit of a two-dimensional map, *g*_*i*_ the goodness-of-fit of the *i*th observation and *g*_*ij*_ the goodness-of-fit of the distance between observations *i* and *j*. After creating an MDS plot as outlined above, the question arises whether the map gives a good approximation to the original distance matrix. If the distance matrix is Euclidean, with **B** having only non-negative eigenvalues, then goodness-of-fit of a *k*-dimensional solution is usually assessed by computing the statistic

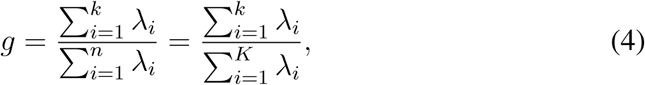

where *K* is the rank of **B**, such that the trailing *n* − *K* zero eigenvalues can be ignored. A common choice is *k* = 2, though in many applications dimensions be-yond the second can be informative. If B has negative eigenvalues, the denominator of Equation (4) is often adjusted by taking absolute values of the eigenvalues, or by considering positive eigenvalues only. Statistic *g* corresponds to the fraction of the total sum-of-squares of the squared distances that is accounted for by the *k*-dimensional approximation, and this fraction can also be obtained as

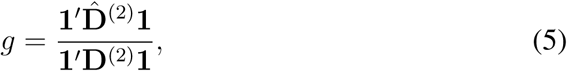

where 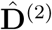 is the approximation obtained by using the first *k* columns of **X** only. The eigenvalues obtained in PCO are proportional to the eigenvalues obtained by a principal component analysis (PCA) of the original data matrix from which the distance matrix **D** has been calculated, in case such a data matrix is available (Mardia et al., 1979) In PCA, these eigenvalues represent the goodness-of-fit of the centred data matrix, leading to the surprising situation that the latter would *equal* the goodness-of-fit of the distance matrix if Equations (4) or (5) are used as the criterion.

### 2.2 Euclidean distance matrices

The total variability in the distance matrix can be expressed as the sum of all squared distances, and this variability is also proportional to the sum of all eigen-values and the trace of **B**. We have

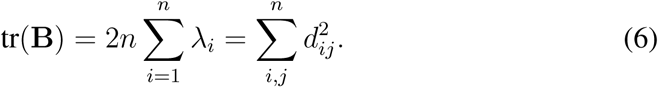

This quantity can be decomposed over dimensions and over observations. This decomposition is given by the *n* × *n* matrix **Q**_*d*_, obtained by

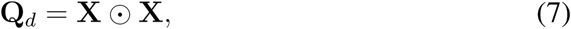

with 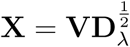, and ⊙ representing the Hadamard product. We use subindex *d* to emphasise this is the decomposition of the total sum of squared distances. Matrix **Q**_*d*_ satisfies

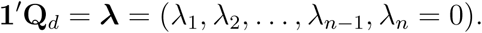

Each row makes a contribution to the total sum-of-squares, and these contributions are given by **w** = **Q**_*d*_**1**. For the *i*th row, this contribution is

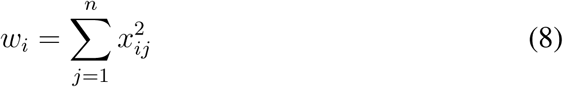

where *x*_*ij*_ represents the entry of row *i* and colum *j* of the solution matrix **X**. The goodness-of-fit for a particular point (*g*_*i*_) in a *k*-dimensional solution can then be calculated as:

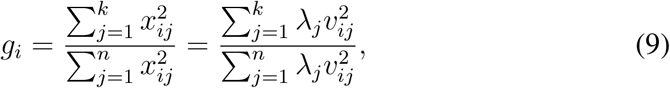

where *v*_*ij*_ represents the *i*th element of eigenvector *j*, the *j*th column of **V**. Statistic *g*_*i*_ indicates how well the contribution of the *i*th row is accounted for in *k* dimensions. At the same time, *g*_*i*_ is the ratio of the squared Euclidean distance between point *i* and the origin in the map and the squared Euclidean distance between point *i* and the origin in the full space. A point will have a large goodness-of-fit if the first *k* eigenvalues are large, and also if it has a large distance from the origin. The overall goodness-of-fit of a *k*-dimensional approximation to the distance matrix is seen to be a weighted average of the goodness-of-fit of each row:

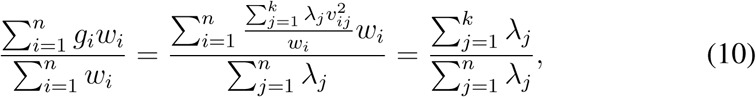

where the weights are the contributions of each row to the total sum-of-squares. Pairwise goodness-of-fit statistics can also be developed. They indicate how well the distance between a particular pair of points is represented. They may be considered more interesting, since our goal is representation of interpoint distances. We use *g*_*ij*_ to refer to the goodness-of-fit of the distance between points *i* and *j*. A natural measure is

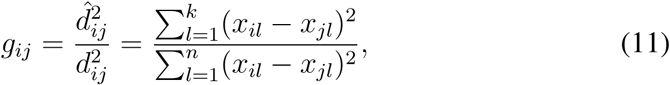

which for Euclidean distance matrices satisfies 0 *≤ g*_*ij*_ *≤* 1.

### 2.3 Non-Euclidean distance matrices

For non-Euclidean distance matrices, a perfect representation of the distance matrix in high-dimensional space is not possible, as is evidenced by the existence of negative eigenvalues in the solution. This complicates the definition of adequate goodness-of-fit measures, both globally, as well as for individual observations and pairs of observations. For an Euclidean distance matrix, the amount of error in a low dimensional approximation will decrease as more dimensions are considered. Finally, an Euclidean distance matrix will be perfectly represented if all non-trivial dimensions with *λ* > 0 are considered. We consider several approaches for obtaining point and pair-wise goodness-of-fit statistics in the non-Euclidean case outlined in the subsections below. We subsequently use an Euclidean subspace, the scalar product matrix, and error statistics for goodness-of-fit calculations.

#### Euclidean subspace

For a non-Euclidean distance matrix, the amount of error in a low dimensional representation will at first decrease as more dimensions are considered, but only up to a certain limiting number of dimensions 𝓁. This number 𝓁 is given by the number of positive eigenvalues that are larger than the absolute value of the last most negative eigenvalue obtained in the analysis. Another particularity is that the coordinates of the solution, as calculated by Equation (3), are only defined for those dimensions that have non-negative eigenvalues. Let *p* be the number of non-negative eigenvalues. We will have *n* eigenvalues, but only *p* coordinates are defined, and only up to 𝓁 coordinates improve the representation of the distance matrix. Let *X*_𝓁_ contain the first 𝓁 coordinates of the solution only. We can compute an estimate of the original distance matrix by computing the Euclidean distances between the observations in 𝓁 dimensions only. One could use the same pointwise goodness-of-fit index as in Equation (9), but using only 𝓁 dimensions in the denominator, and using at most *k ≤* 𝓁 dimensions.

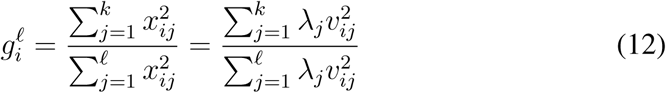

A pairwise measure, analogous to Equation (11), is

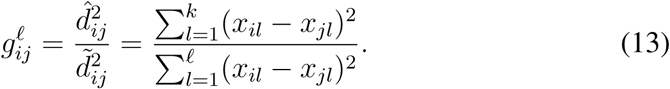

These measures assume that 𝓁 ≥ 2.

#### Scalar product matrix

Mardia (1978; 1979) has suggested the use of the *squared* eigenvalues for non-Euclidean distance matrices. Because the distances are approximated indirectly, via the scalar product matrix **B**, one could report the goodness-of-fit of the latter instead. The total sum-of-squares of **B** is given by tr 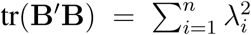, and therefore the goodness-of-fit of **B** is obtained by using the *squared* eigenvalues

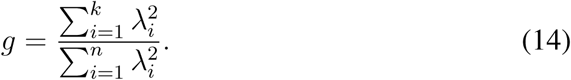

Akin to Equation (7), we now have the decomposition

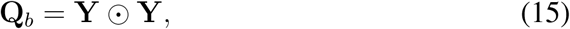

with **Y** = **VD**_*λ*_. We use subindex *b* to emphasize this is the decomposition of the total sum of squares of the scalar product matrix **B**. Matrix **Q**_*b*_ satisfies

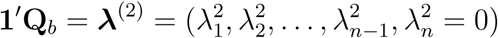

Contributions of each row to the total sum-of-squares of **B** can be obtained as

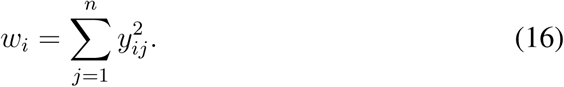

The goodness-of-fit for a particular point (*g*_*i*_) in the *k*-dimensional solution can then be calculated as:

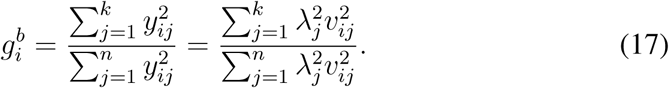

As before, a weighted average of these point-wise measures gives the overall goodness-of-fit. It is harder to develop a useful measure for the goodness-of-fit of pairwise distances in the non-Euclidean case. For Euclidean distance matrices, the approximation to the observed distances is always from below, and therefore Equation (11) seems a sensible measure, with an upper bound of 1 if the distance of the corresponding pair is perfectly represented. In the non-Euclidean case, the fitted distances can exceed the originally observed distances (see the geographical example in the next section).

#### Error statistics

Instead of focusing on goodness-of-fit, one can also focus on error. If the map obtained by MDS is good approximation, then errors will be small, and concentrated around zero. Most points may be expected to make a small and similar contribution to the error sum-of-squares (ESS). A simple measure of poorness-of-fit, indicated by 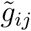, is the contribution to the total ESS,

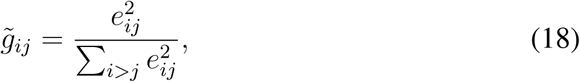

where *e*_*ij*_ is an element of 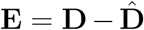. Large outliers on 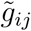 correspond to poorly fitted pairs. These error contributions satisfy 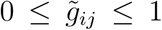. Equation (18) can also be used to develop a pointwise statistic, by summing errors that pertain to a particular observation, that is

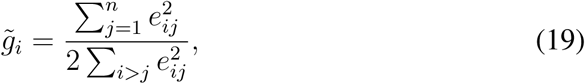

and large outliers on 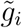 would correspond to poorly fitted observations. By counting each error twice in the denominator, we achieve that 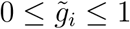.

## 3 Examples

We illustrate the proposed goodness-of-fit statistics with three different data sets taken from demography, genetics and geography and discuss these in the following subsections.

### 3.1 Demographic distances between countries

We discuss the analysis of an Euclidean distance matrix obtained from a demographic dataset of six variables (live birth rate, death rate, infant death rate, male and female life expectancy and gross national product (GNP)) for 97 countries described by Rouncefield (1995). Euclidean distances between countries were calculated using the standardized data. GNP was log-transformed to linearize its relationship with the other variables prior to standardization. An MDS map of the Euclidean distance matrix of the countries is shown in Figure 1A; The original variables have been mapped into the MDS plot by regression to aid interpretation (Graffelman and Aluja-Banet, 2003). This shows the first dimension is a wealth dimension separating rich countries with high life expectancies on the left from poor countries with high infant death rates and high birth rates on the right. In Figure 1B the countries are colour coded according to their goodness-of-fit, and this reveals some countries, mainly in the center of the map, with, according to Equation (9) a low goodness-of-fit (Saudi Arabia 0.26; Libya 0.29; Oman 0.29). These points are maybe better ignored (or brushed away) to avoid misinterpretation of the map. In Figure 1C we use pairwise goodness-of-fit statistics to reveal relatively poorly displayed inter-country distances that have *g*_*ij*_ < 0.50; these countries are connected by dotted lines. This reveals that many countries that appear as neighbours in the map are in reality father away from each other than the map suggests, being their squared full space distance at least two times larger. In Figure 1D we focus on well-represented inter-country distances, showing all distances for Sierra Leone that have *g*_*ij*_ > 0.90. This shows almost all distances with respect to Sierra Leone are very well represented, and that the country acts as an *attractor point*. Similarly, Gambia, Malawi, Ethiopia, Somalia, Angola and Mexico are also well represented attractor points in terms of inter-country distances. These countries have, in terms of the original distance matrix, a large average distance with respect to the rest of the countries. In this analysis, the larger distances have in general, a better fit than the smaller distances, as is also revealed by a scatter plot of observed against fitted distances (See supplementary Figure S1).

**Figure 1:**
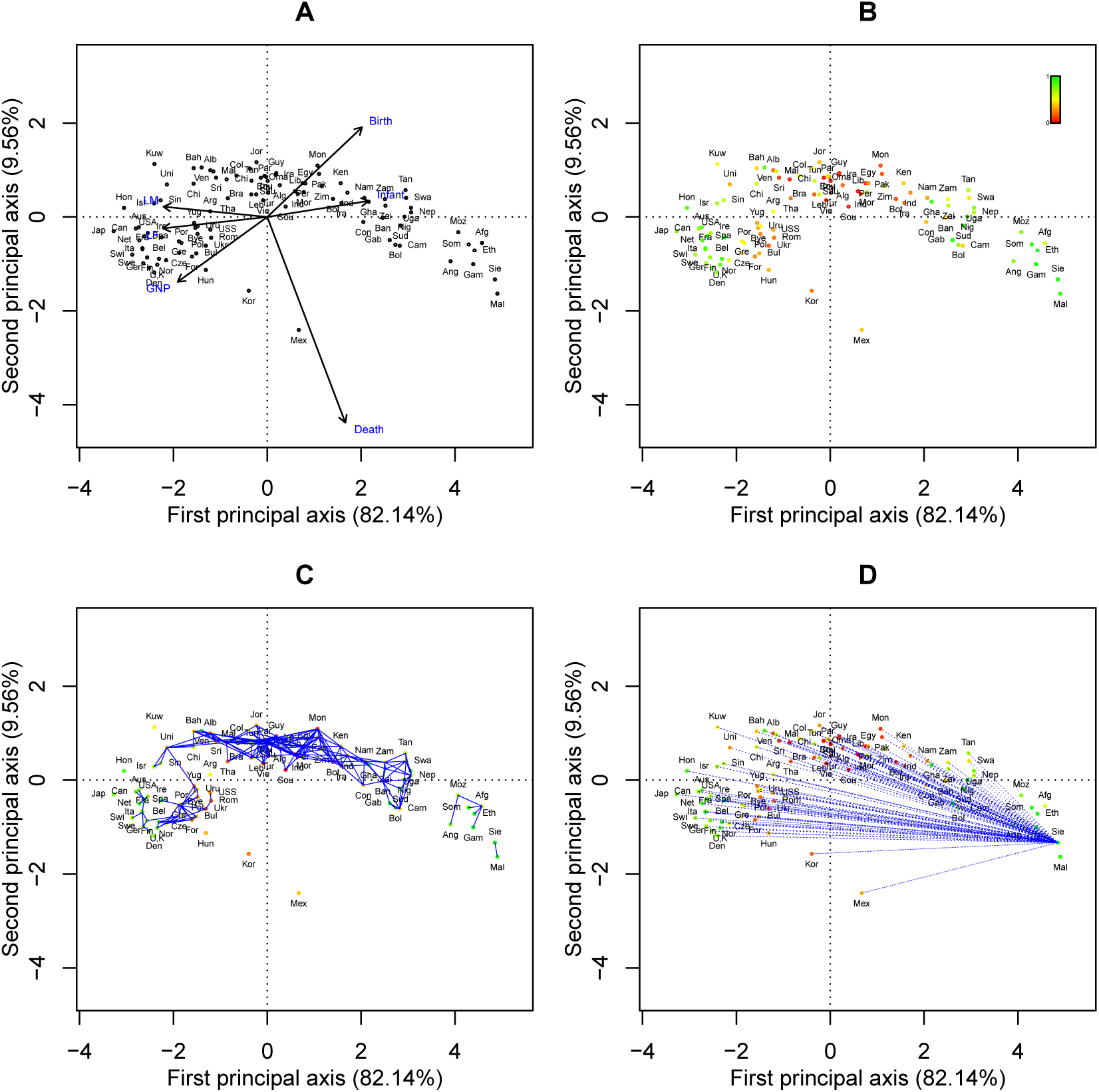
MDS of the poverty data set. A: MDS map with added variables (LM: life expectancy of males, LF: life expectancy of females, GNP: gross national product, Birth: birth rate, Infant: infant death rate, Death: death rate). B: MDS map colour-coded according to the goodness-of-fit of the countries. C: MDS map showing inter-country distances with fit below 0.50. D: MDS map showing inter-country distances with fit above 0.90 for Sierra Leone.

### 3.2 Genetics distances between individuals

MDS is widely used in genetics for the detection of population substructure, which refers to the existence of groups of individuals in a genetic database that come from different human populations. Many examples of the use of MDS for this purpose can be found in the genetic literature (Pemberton et al., 2010; Jakobsson et al., 2008; Sabatti et al., 2009; Wang et al., 2010; Pemberton et al., 2013). MDS studies in genetics often use the allele sharing distance. The possible genotypes of bi-allelic genetic variables are, in generic notation, the homozygotes AA and BB and the heterozygote AB. If one of the alleles is counted, usually the minor allele, the genotype data can be coded into 0,1,2 format. The allele sharing distance is defined as

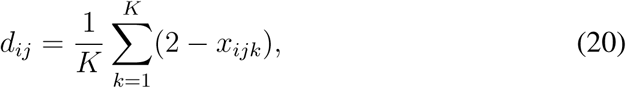

where *x*_*ijk*_ is the number of alleles shared by individuals *i* and *j* at genetic variable *k*, taking only values in the set (0,1,2). Equation (20), when applied to (0,1,2) data, is actually equivalent to the Manhattan distance, which is a well-known metric in the statistical literature (Mardia et al., 1979). This metric is often also referred to as the city-block distance or the taxicab metric. The Manhattan distance is directly proportional to the allele sharing distance. Geneticist often perform MDS of genetic data with the PLINK software (Purcell et al., 2007); this program does classical metric MDS as described above on the Manhattan distances between the individuals of a genetic database. We present here an MDS of 109 initially presumed unrelated individuals of a sample of Chinese in Metropolitan Denver, CO, USA; the CHD sample of the 1000 Genomes project (www.internationalgenome.org).

Genetic variants were filtered prior to MDS by selecting only autosomal variants without missing values, with a minor allele frequency above 0.40, non-significant in an exact test for Hardy-Weinberg equilibrium (*α* = 0.05), and without strong correlations with flanking markers. These filters left 28.158 genetic variables in the analysis. Figure 2A shows the MDS map of the individuals. All eigenvalues obtained were non-negative. The overall goodness-of-fit for a two-dimensional display is low, only 3.0%, which is typical of genetic applications with large datasets. Figure 2B applies a goodness-of-fit filter of 0.25 at the individual level. This reveals that most indvididuals have a poor fit, and only the four outliers are well represented. The four outliers correspond to individuals for which a family relationship has been identified by Pemberton et al. (2010). In particular, the outlying pair in the first dimension has been estimated to be a full sib (FS) pair (identifiers NA17981 and NA17986), whereas the outlier in the second dimension has been estimated to be a parent-offspring (PO) pair (identifiers NA17976 and NA18166). Figure 2C connects individuals with a poorly fitted genetic distance (goodness-of-fit < 0.25), whereas Figure 2D connects individuals with relatively better represented genetic distances (goodness-of-fit > 0.50). This reveals that most genetic distances between unrelated individuals around the origin are poorly displayed, and that only the genetic distances of the related individuals are reasonably well fitted. Supplementary Figure S2 shows the scatterplot of observed against fitted Manhattan distances, and also confirms that all distances involving one or more individuals that form part of a the single PO or FS family relationship are better fitted. Interestingly, the third dimension of the MDS map shows another pair of outliers, corresponding to a second degree relationship that was also uncovered by Pemberton et al. (2010).

**Figure 2:**
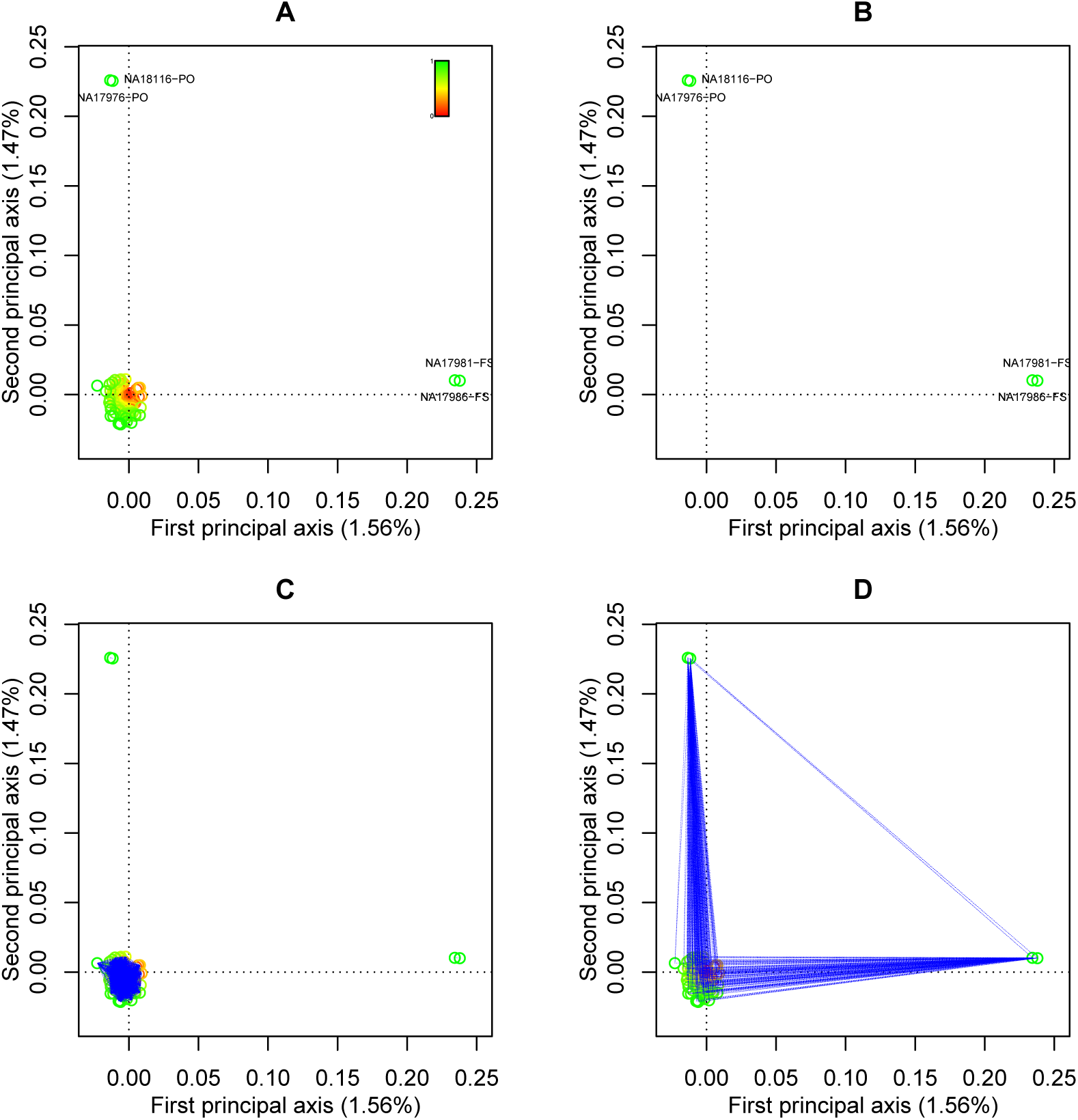
MDS of CHD data set. Individuals are colour-coded according to their goodness-of-fit. A: MDS of the Manhattan distances. B: Filtered MDS map where individuals with goodness-of-fit below 0.25 are not shown. C: MDS map where individuals with a pairwise goodness-of-fit below 0.25 are connected by blue lines. D: MDS map where individuals with a pairwise goodness-of-fit above 0.50 are connected by blue lines.

### 3.3 Geographic distances between Spanish cities

A classical example of metric MDS is the construction of a geographical map of a set of cities based on a table of inter-city distances. Such distances can be given in different form: as straight line geographical distances, road or railway distances, or as travel times. Many examples of this kind have been described in the literature (Mardia et al., 1979; Manly, 1989; Johnson and Wichern, 2002). We consider the road distances in kilometers between 47 cities in Spain. Figure 3A shows the result of a classical metric MDS of the data. The first principal axis is shown in the vertical dimension in order to better match the geographical map of Spain. There are 24 positive eigenvalues, the 25th eigenvalue is the structural zero, and the remaining 22 are negative. Of all positive eigenvalues, three exceed the largest negative eigenvalue in absolute value. Using the standard adjustments, the goodness-of-fit of the distance matrix in two dimensions is 0.699 (using absolute values) or 0.815 (using positive eigenvalues only). The goodness-of-fit of the scalar products, obtained by using squared eigenvalues, is 0.979. Symbols in Figure 3A are colour coded to indicate goodness-of-fit of the observations according to Equation (17), which is based on squared eigenvalues. This suggests that the more outlying peripheral cities in Catalonia and Andalusia have a better fit than the Bask cities in the north and the central cities. The most central cities, Guadalajara and Madrid, have the poorest goodness-of-fit, 0.23 and 0.45 respectively. Figure 3B shows boxplots of the pointwise statistics of equations Equations (17) (left) and (12) (right). Both statistics suggest about three relatively poorly fitted cities: Madrid, Cuenca and Guadalajara. Figure 3C shows these cities have a lower average distance with respect to the other Spanish cities, and that cities with larger average distances tend to have better fit. Figure 3D shows the pairs of cities that contribute more than 1% to the total error sum-of-squares, according to Equation (18); the distance between Guadalajara and Cuenca has the worst fit. A scatter plot of fitted against observed distances is shown in supplementary Figure S3, and shows graphically, despite the negative eigenvalues, an excellent fit with only a few poorly fitted intercity distances. The outliers in Figure S3 effectively correspond to the distance traced by lines in Figure 3D.

**Figure 3:**
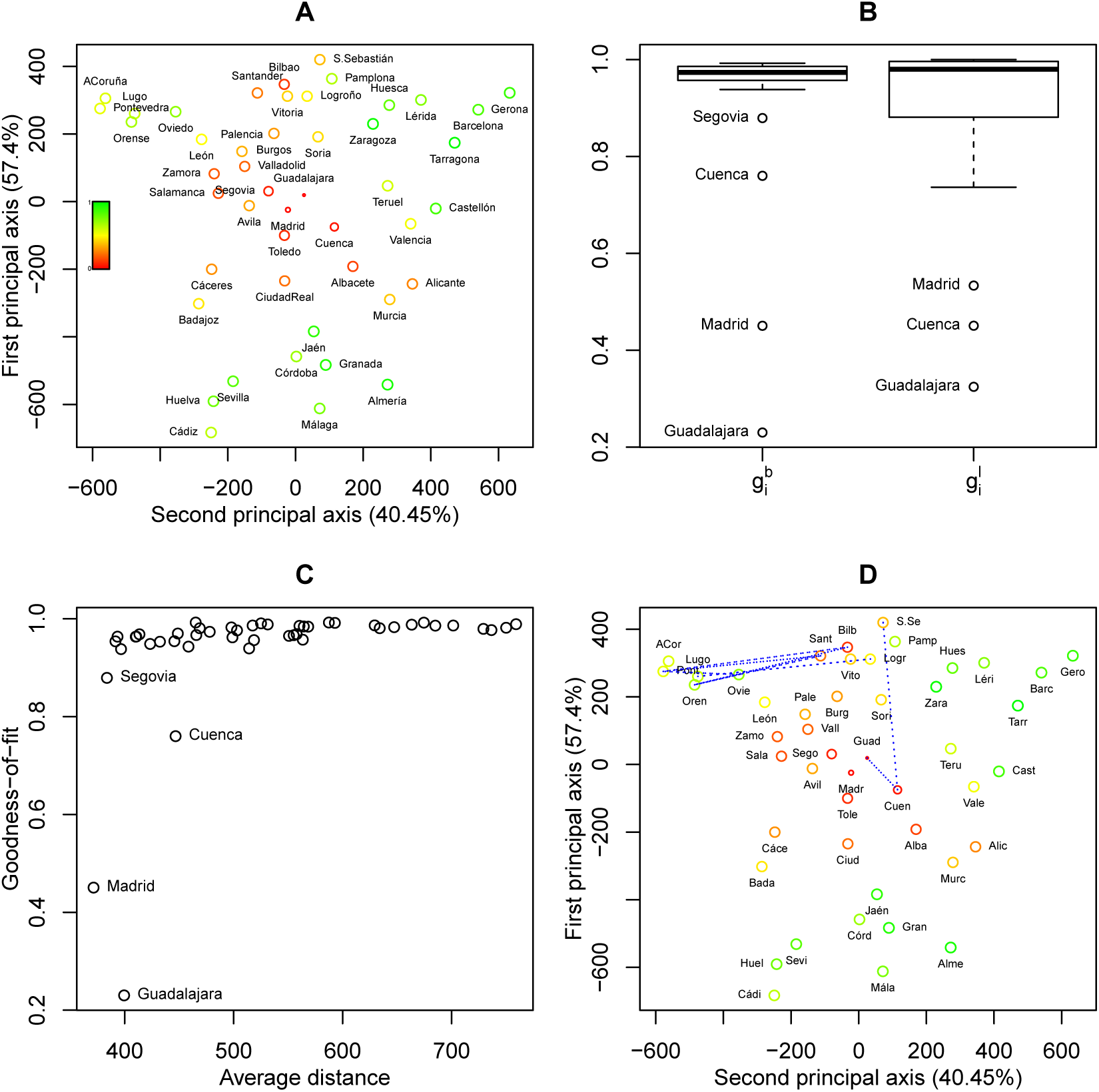
Metric MDS of intercity distances in Spain. A: MDS map with cities colour-coded and symbols scaled according to the goodness-of-fit of the city. B: Boxplots of pointwise goodness-of-fit measures according to Equations (17) and (12)). C: Goodness-of-fit 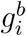 as a function of the average distance. D: MDS map with poorly represented distances, as identified by Equation (18) with 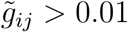, connected by blue lines.

## 4 Discussion

We have developed statistics that quantify the goodness-of-fit of observations and of pairs of observations in classical metric MDS. Nowadays, MDS is applied to increasingly large datasets, and the proposed statistics can be used to identify poorly represented (pairs of) points. Such points may be brushed away in order to emphasise the most salient aspects of the analysis and to avoid misinterpretations. We stress that the brushing of a point is not the same as its elimination from the analysis. A brushed point has been used in the analysis, but is simply not shown because it is poorly fitted. We do not suggest a re-analysis of the data without the poorly fitted points, as this would give an entirely new map, where again poorly fitted observations can be expected to be present. The practical applications in this paper show that poorly represented observations often cluster around the origin of the MDS map. However, goodness-of-fit filtering in MDS is not equivalent to brushing away all points within a circle around the origin. Indeed, a point close to the origin can be well represented if its distance from the origin in the full space is also small.

In many applications of classical metric MDS, negative eigenvalues arise. Overall goodness-of-fit is then usually expressed by an ad hoc adjustment, e.g. considering only positive eigenvalues, or taking absolute values of the eigenvalues. These adjustments do not have a sound theoretical foundation, and are used merely to avoid that the good-of-fit statistic in Equation (4) exceeds one. When the squared eigenvalues (14) are used, no ad-hoc adjustments are needed, as the index is always neatly in the 0-1 range. The use of squared eigenvalues implies we report the goodness-of-fit of **B** instead of the goodness-of-fit of the distance matrix. Because the distances are approximated indirectly, via scalar product matrix **B**, it is then understood that a better fit of **B** will generally imply a better fit of **D**, without quantifying exactly how well **D** is actually represented. A disadvantage is that, beyond 𝓁 dimensions, the goodness-of-fit of **B** increases as more dimensions are included, while the goodness-of-fit of **D** is actually deteriorating. Overall goodness-of-fit statistics will of course look more favourable if squared eigenvalues are used.

An attractive property of classical metric MDS is that it provides a solution that is in the same scale as the original distance matrix. If the original distance matrix is in kilometers, the MDS map is also in kilometers, which facilitates interpretation of the map. This is obvious for the geographical data set analysed above, but also for the genetic data, as described next. For genetic data coded in (0,1,2) format, it is convenient to scale the Manhattan distance matrix by 1*/k* (or, accordingly, scale by 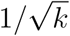 if Euclidean distances are used). This scaling will not affect the configuration of the points in the map, and neither its goodness-of-fit, but it will render the axes and distances of the map more interpretable. Two individuals that are a unit distance apart in the map now differ on average by one allele per locus. This scaling will typically bring all the coordinates in the MDS map within the (-1,1) interval, as the maximum difference in number of alleles between two individuals is two (see Figure 2). This interpretation is hampered by the fact that the map is a low-dimensional approximation to the original distance matrix, and therefore the property will not hold exactly, but only approximately so. For distance matrices that have the Euclidean property, and that therefore only have non-negative eigen-values (Mardia et al., 1979, Chapter 14), the map distances will approximate the true distances from below, and one can say that two individuals that are one unit apart, will differ on average by at least one minor allele per locus. The ability of classical MDS to detect related individuals arises from it sensitivity to outliers: the related individuals have the smallest possible distance in the whole database (see Figure S2), and the plane fitted by MDS is tilted toward these individuals. Consequently, all distances involving these outliers are relatively better represented. At first sight, the MDS map may suggest the outliers to be different from the rest of the sample, potentially stemming from a different population. This is not the case, and in fact the original distances of the outlying individuals with respect to the other individuals of the sample are not larger than the average original distance between just any two unrelated individuals.

## 5 Software and datasets

The function `cmdscale` of the statistical environment R (R Development Core Team, 2004), actually version 3.5.1, is widely used to perform classical multidimensional scaling. This function does not allow the calculation of all statistics proposed in Section 2. We supply R code (function `PrinCoor`) that implements all goodness-of-fit statistics discussed in this paper as supplementary material. All datasets used in this paper are accessible online. The poverty data set is available at http://jse.amstat.org/v3n2/datasets.rouncefield.html, the genetic data set is available at http://www.internationalgenome.org and the geographical data set is available at http://www-eio.upc.es/~jan/data/SpainDist.dat.

## 6 Acknowledgements

This work was partially supported by grants RTI2018-095518-B-C22 of the Spanish Ministry of Science, Innovation and Universities and the European Regional Development Fund, and by grant R01 GM075091 from the United States National Institutes of Health.

## 7 Supplementary figures

**Figure 4:**
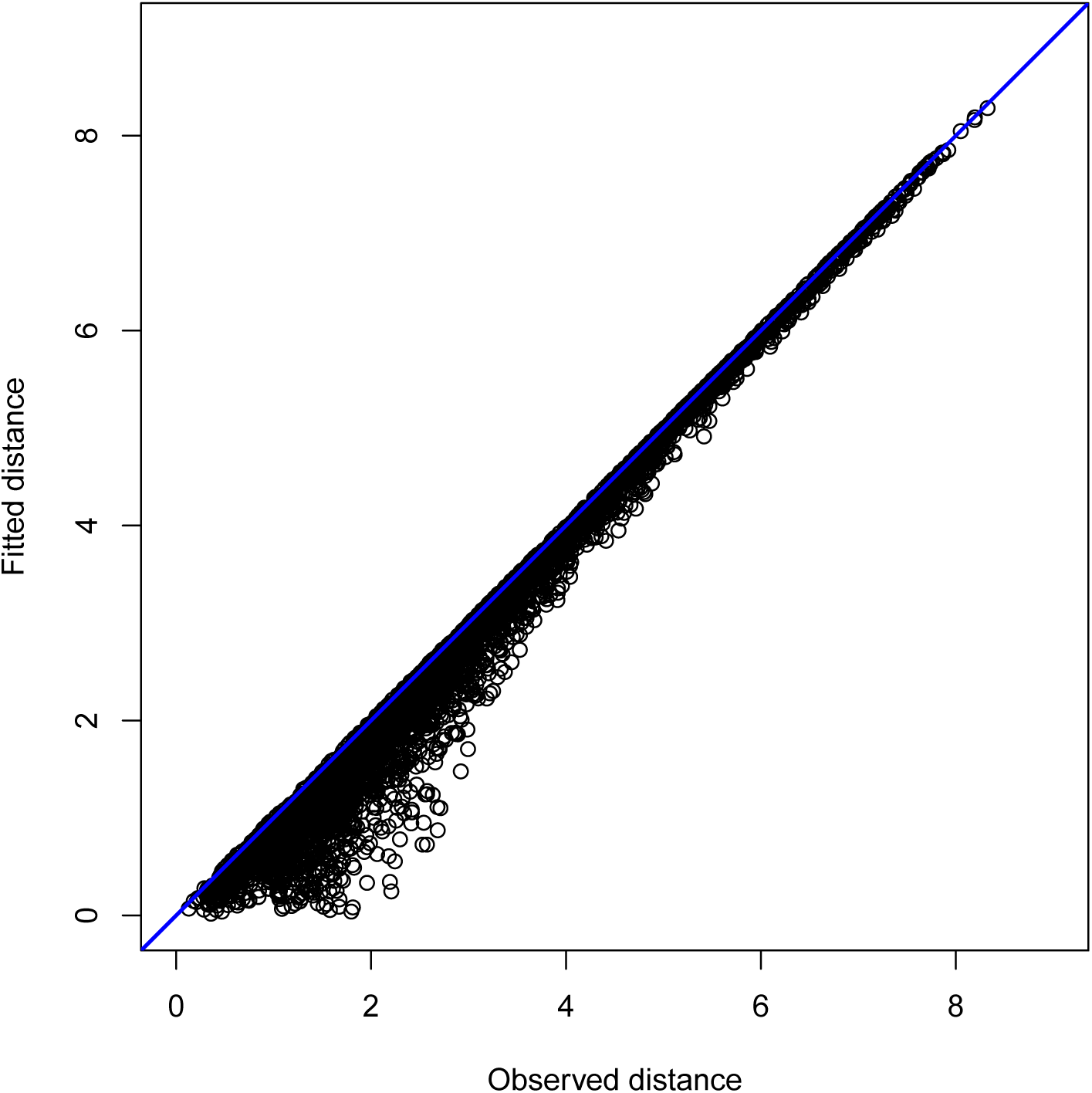
Poverty data set. Plot of fitted against observed distances.

**Figure 5:**
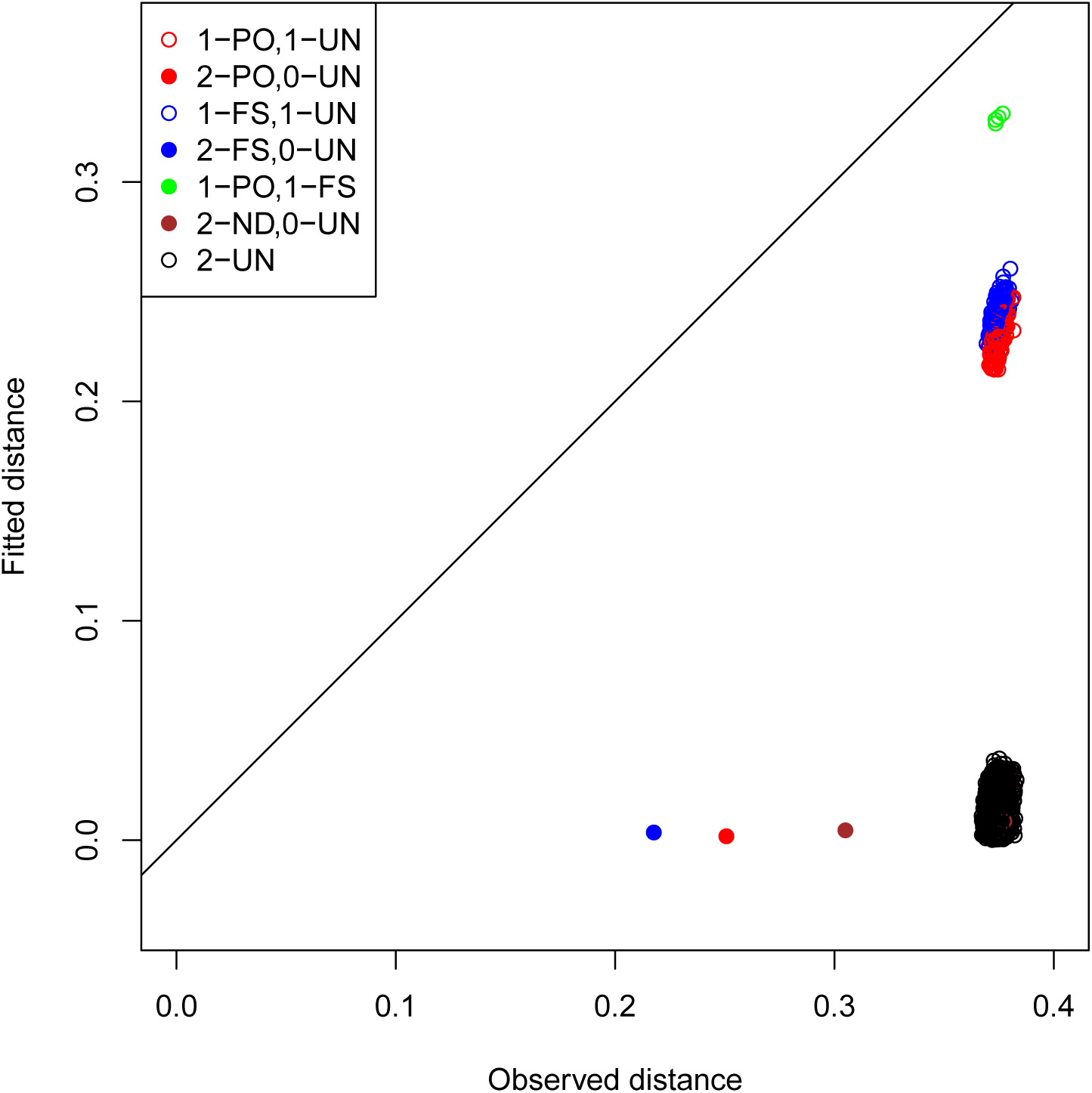
CHD data set. Plot of fitted against observed distances. Pairs are coloured according to the number of individuals that participate in the single FS and PO relationships in the dataset.

**Figure 6:**
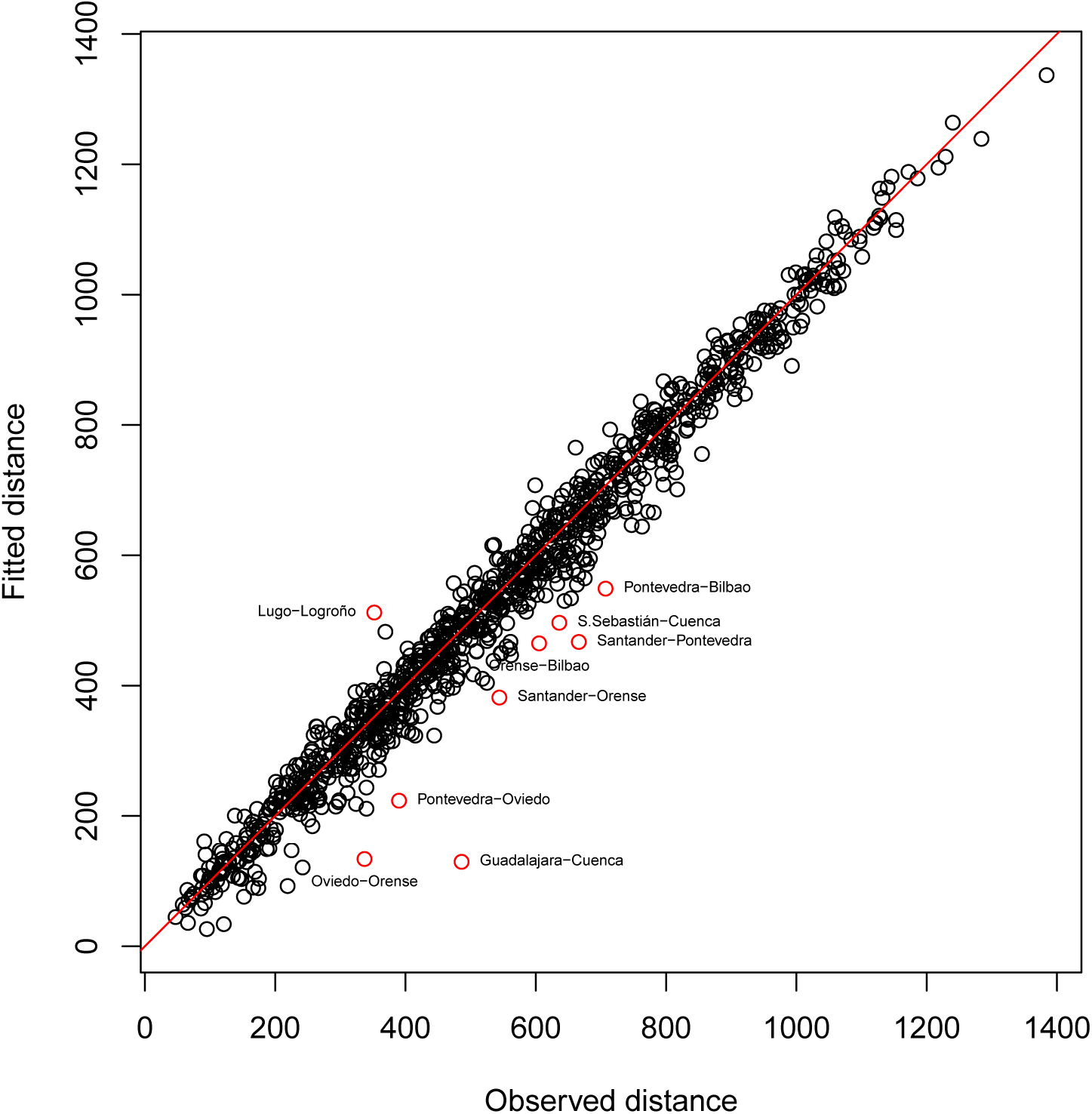
Spanish distances data set. Plot of fitted against observed distances. Poorly fitted points are marked in red.

## References

Borg, I. and Groenen, P. J. F. 2005. Modern Multidimensional Scaling. Springer, New York, second edition.

Carroll, J. and Chang, J. 1970. Analysis of individual differences in multidimensional scaling via an n-way generalization of Eckart-Young decomposition. Psychometrika, 35(3):283–319.

Cox, T. F. and Cox, M. A. A. 2001. Multidimensional scaling. Chapman & Hall, second edition.

De Leeuw, J. and Mair, P. 2009. Multidimensional scaling using majorization: SMACOF in R. Journal of Statistical Software, 31(3).

Gower, J. C. 1966. Some distance properties of latent root and vector methods used in multivariate analysis. Biometrika, 53(3):325–338.

Graffelman, J. and Aluja-Banet, T. 2003. Optimal representation of supplementary variables in biplots from principal component analysis and correspondence analysis. Biometrical Journal, 45(4):491–509.

Jakobsson, M., Scholz, S., Scheet, P., Gibbs, J., VanLiere, J., Fung, H., Szpiech, Z., Degnan, J., Wang, K., Guerreiro, R., Bras, J., Schymick, J., Hernandez, D., Traynor, B., Simon-Sanchez, J., Matarin, M., Britton, A., van de Leemput, J., Rafferty, I., Bucan, M., Cann, H., Hardy, J., Rosenberg, N., and Singleton, A. 2008. Genotype, haplotype and copy-number variation in worldwide human populations. Nature, 451:998–1003.

Johnson, R. A. and Wichern, D. W. 2002. Applied Multivariate Statistical Analysis. New Jersey: Prentice Hall, fifth edition.

Kruskal, J. 1964. Multidimensional scaling by optimizing goodness of fit to a nonmetric hypothesis. Psychometrika, 29:1–27.

Manly, B. F. J. 1989. Multivariate statistical methods: a primer. Chapman and Hall, London.

Mardia, K. V. 1978. Some properties of classical multi-dimensional scaling. Communications in Statistics – Theory and Methods, 7(13):1233–1241.

Mardia, K. V., Kent, J. T., and Bibby, J. M. 1979. Multivariate Analysis. Academic Press London.

Pemberton, T., DeGiorgio, M., and Rosenberg, N. 2013. Population structure in a comprehensive genomic data set on human microsatellite variation. G3: Genes, Genomes, Genetics, 3(5):891–907.

Pemberton, T., Wang, C., Li, J., and Rosenberg, N. 2010. Inference of unexpected genetic relatedness among individuals in HapMap Phase III. American Journal of Human Genetics, 87:457–464.

Purcell, S., Neale, B., Todd-Brown, K., Thomas, L., Ferreira, M., Bender, D., Maller, J., Sklar, P., de Bakker, P., Daly, M., and Sham, P. 2007. Plink: A toolset for whole-genome association and population-based linkage analysis. American Journal of Human Genetics, 81(3):559–575.

R Development Core Team 2004. R: A language and environment for statistical computing. R Foundation for Statistical Computing, Vienna, Austria. ISBN 3-900051-00-3.

Rouncefield, M. 1995. The statistics of poverty and inequality. Journal of Statistics Education, 3(2):null.

Sabatti, C., Service, S., Hartikainen, A., Pouta, A., Ripatti, S., Brodsky, J., Jones, C., Zaitlen, N., Varilo, T., Kaakinen, M., Sovio, U., Ruokonen, A., Laitinen, J., Jakkula, E., Coin, L., Hoggart, C., Collins, A., Turunen, H., Gabriel, S., Elliot, P., McCarthy, M., Daly, M., Järvelin, M., Freimer, N., and Peltonen, L. 2009. Genome-wide association analysis of metabolic traits in a birth cohort from a founder population. Nature genetics, 41(1):35–46.

Torgerson, W. 1958. Theory and Methods of Scaling. Wiley, New York.

Wang, C., Szpiech, Z., Degnan, J., Jakobsson, M., Pemberton, T., Hardy, J., Singleton, A., and Rosenberg, N. 2010. Comparing spatial maps of human population-genetic variation using procrustes analysis. Statistical applications in genetics and molecular biology, 9. Article 13.

